# “Low-cost” initial burst of root development in whole *Fagus crenata* seedlings: The key to survival?

**DOI:** 10.1101/535500

**Authors:** Yoko Kurosawa, Shigeta Mori, Mofei Wang, Juan Pedro Ferrio, Keiko Yamaji, Kenichi Yoshimura, Citra Gilang Qur’ani

## Abstract

Terrestrial plants are rooted in one place, and therefore their metabolism must be flexible to adapt to continuously changing environments. This flexibility is probably influenced by the divergent metabolic traits of plant organs. However, direct measurements on organ-specific metabolic rates are particularly scarce and little is known about their roles in determining whole-individual meatabolism. To reveal this on seedlings of *Fagus crenata*, which is one of the most widespread dominant genus in temperate deciduous broad leaf forests in the circum-polar Northern Hemisphere, we measured respiration, fresh mass and surface area for total leaves, stems and roots of 55 individuals in two years from germination and analyzed their relationships with individual metabolism. Proportion of roots to whole plant in mass increased from approximately 17% to 74%, and that in surface area increased from about 11% to 82% in the two years. Nonetheless, the increment of the proportion of root respiration to whole-plant respiration was from 9.2% to only 40%, revealing that the increment in mass and surface area of roots was much larger than the increment in energetic cost. As a result, only the roots showed a substantial decline in both respiration/surface area and respiration/mass among the three organs; roots had about 90% decline in their respiration/surface area, and 84% decline in their respiration/mass, while those in leaves and stems were relatively constant. The low-cost and rapid root development is specific to the two years after germination and would be effective for avoiding water and nutrient deficit, and possibly helps seedling survival. This drastic shift in structure and function with efficient energy use in developmental change from seeds to seedlings may underpin the establishment of *F. crenata* forests. We discuss significance of lowering energetic cost for various individual organisms to effectively acquire resources from a wide perspective of view.

## Introduction

Individual metabolism is a fundamental process that transforms energy and materials to support various biological processes as a base for adaptation to changing environments [1,2]. Therefore, the metabolic rate has profound physiological, ecological and evolutionary implications [3,4], which would be a key to understand and predict the effects of climate change on organisms and ecosystems [1,2].

In general, the metabolic rate (i.e. respiration rate, *R*) of individual organisms scales with body size (*X*), and is usually described as the simple power function of body mass:

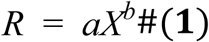

where *a* is a normalization constant, and *b* is the scaling exponent (slope on the log-log coordinates) [5–7]. The equation (1) represents the emergent outcomes of the metabolism of individuals under various constraints [2,8,9]. Therefore, to obtain a mechanistic insight into the regulation of scaling of metabolic rate, we need empirical evidence of whole-organism measurements. However, little is known about the relationships between metabolic rate and body size with the reliable data from small to giant individuals [10]. This is because most of the studies on metabolic scaling have been based on indirect evidence and aimed to construct theoretical models to explain the exponent *b*, which has widely been assumed to be 3/4 as suggested by the WBE model [5–9]. The size scaling of individual organisms results from the sum of differentiated organs with distinctive functions and structure, each one showing contrasting responses under changing environments [11–13]. Therefore, evaluating metabolic rate of each organ is crucial to understand the scaling of metabolism associating with body size, namely mass or surface area.

Terrestrial plants are supposed to adapt to various environments by adjusting the biomass partitioning among organs, as typically shown in the root/shoot ratio [11–17]. To date, the optimal partitioning theory has mainly evaluated allocation between shoot and roots in mass. It suggests that plants should allocate more biomass to the shoot when limiting resource is carbon and to the roots when limiting resource is water or nutrient [11,17]. However, in spite of the significant implication of metabolic rate, few studies have compared metabolic rate of shoot (leaves + stems) and roots at the whole-plant level [18]. The comparison of respiration rate between shoot and roots, with measurement of organ-specific respiration at the individual level, would provide a new insight into the energy partitioning and would progress our understanding about whole-plant adaptation for resource acquisition.

The purpose of this study is to understand the processes of establishment of individual seedlings of *Fagus crenata* under varying environments on the basis of the partitioning theory [11,17], considering the allocation of individual mass, surface area, and energy to the different organs. *Fagus* is one of the most widespread dominant genus in temperate deciduous broad leaf forests in the circum-polar Northern Hemisphere [19–21]. Under low light conditions, *Fagus* seedlings are characterized by high mortality in current year of germination, which is called as bottleneck phase [22–26]. We hypothesize that the effective adaptation of seedlings for survival beyond the bottleneck phase is largely dependent on the partitioning of energy within each individual. To test this assumption, we need size-scaling of respiration that cover seedlings in current-year of germination and 1-year old.

Here, we show the respiration rates of total leaves, stems, and roots of various sized seedlings and size-scaling of organ-specific respiration to discuss the shift of the partitioning of energy within individuals during the two years after germination. To date, little is known about the whole-plant physiology of *Fagus* seedlings in the bottleneck phase. Our empirical study on the partitioning of energy would clarify how seedlings survive over the bottleneck phase, which would greatly affect the population dynamics of forest in the circum-polar Northern Hemisphere under changing environment [14,26,27]. Finally, we discuss efficient energy use in divergent individual organisms to effectively acquire resources.

## Materials and methods

### Ethics statement

Our study included fieldwork activities for collecting *F. crenata* seeds and seedlings, and were conducted in Japanese National Forest. The field work was permitted by the Shonai District Forest Office and did not involve any endangered or protected species.

### Seed collection and plant materials

The measurements were performed on current-year (n = 46) and 1-year-old (n = 9) seedlings of *F. crenata*, raised from seeds in pots outside the Yamagata University Tsuruoka Campus (38°73′N, 139°82′E). The seeds were collected from a mature forest of *F. crenata* in Tsuruoka (38°30′N, 139°57′E), Yamagata prefecture in 2015 and 2016 to prepare 1-year-old and current-year seedlings, respectively.

The pots were filled with commercially available Kanuma pumice mixed with leaf mold, and kept in a place with sufficient natural light and well-watered. We conducted every measurement at the whole-plant level, using total leaves (including cotyledon), stems, and roots. The whole-plant fresh mass ranged between 41.2×10^−5^ and 23.5×10^−3^ (kg) from the smallest current-year seedling to the largest 1-year-old individual, as compiled in Table 1.

**Table 1.** Compilation of the minimum and maximum values of fresh mass, surface area, and respiration of whole plant, roots, leaves, and stems at the individual level among all seedlings (n = 55).

| | Fresh mass<br>($\times 10^{-5}$ kg) | | | Surface area<br>($\times 10^{-4}$ m<sup>2</sup>) | | | Respiration rate<br>($\times 10^{-4}$ μmol sec<sup>-1</sup>) | | |
| --- | --- | --- | --- | --- | --- | --- | --- | --- | --- |
|  | min. | max. | max./min. | min. | max. | max./min. | min. | max. | max./min. |
| Whole plant | 41.2 | 2350 | 57.0 | 23.6 | 1210 | 51.3 | 8.70 | 131 | 15.1 |
| Roots | 7.04 | 1730 | 246 | 2.54 | 998 | 393 | 0.800 | 52.6 | 65.8 |
| Leaves | 19.3 | 172 | 8.91 | 19.2 | 185 | 9.64 | 4.60 | 53.3 | 11.6 |
| Stems | 8.52 | 480 | 56.3 | 1.43 | 31.8 | 22.2 | 1.10 | 42.3 | 38.5 |

Every measurement is including both current-year and 1-year-old seedlings.

### Measurement of respiration rates and surface area

We separated the seedlings into leaves, stems, and roots, and enclosed them separately in custom-made chambers (80 or 160 cm^3^), promoting air circulation within the chamber using a DC fan. We confirmed that the separation did not have an effect on the measured values of whole-plant respiration, as reported by Mori et al. [10]. Increment rates of CO_2_ concentrations within the closed air-circulation system were measured every 5 seconds using an infrared CO_2_ analyzer (GMP343, Vaisala, Helsinki, Finland), and normalized to 20°C assuming a standard Q_10_ = 2. During the measurements, we kept the plant materials wrapped in wet paper to prevent transpiration, and the measurements were taken within 20 minutes of the excavation for each seedling.

Leaf surface area was measured with an area meter (LI-3100C, LICOR, Lincoln, NE, USA), and stem surface area was determined as sum of the surface area of stem sections, following a cylindrical approximation from their diameter and length. Roots were scanned at 800 dpi resolution with a flatbed scanner (Epson Perfection V800, Seiko Epson, Japan), followed by the measurements of root surface area with image analytical software (WinRhizo, Regent Instruments, Quebec, Canada).

### Data analysis

We fitted the respiration–fresh mass and respiration–area scaling relationships using a simple-power function on log-log coordinates, based on reduced major axis regression (RMA) [28] of the log transformed version of equation (1), using all the measurement data of the 55 seedlings from current-year to 1-year-old. We also analyzed the size-scaling values for surface area in relation to fresh mass (based on RMA) for organs, and their respiration per unit mass and surface area in relation to whole-plant mass (based on ordinary least squares regression, OLS) on the log-log coordinates. In the analysis for the size-scaling of respiration per unit mass and surface area, OLS analysis performed adequately rather than RMA. All of the data used for the regression analysis were compiled in S1 File.

## Results

### Whole-plant respiration

Whole-plant respiration reflects individual adaptation as an integrated use of energy partitioned to leaves, stems, and roots. To consider the total energy use of individuals, we showed relationships of whole-plant respiration (*R*; μmol s^−1^) to fresh mass (*M*; kg) and surface area (*S*; m^2^) of the seedlings from current-year to 1-year old on log-log coordinates. We fitted them with a simple power function (1) (Fig. 1, Table 2). Astonishingly, there was no significant difference between the exponents for the scaling of *R* to *M* (*b* = 0.647, 95% CI of *b*: 0.584 to 0.702, r^2^ = 0.898; Fig.1a) and that of *R* to *S* (*b* = 0.685, 95% CI of *b*: 0.625 to 0.738, r^2^ = 0.892; Fig. 1b). The *b* = 0.647 for *R* to *M* was closer to 2/3 ≈ 0.667, being significantly different from the *b* = 3/4 = 0.75 that was predicted by the WBE model.

**Table 2.** Scaling of respiration rate (µmol sec^−1^) of whole plant, roots, leaves, and stems with their fresh mass (kg) and surface area (m^2^) fitted by equation (1).

| Independent variable | | Exponent<br>($b$) | 95% CI of $b$ | Normalization<br>constant ($a$) | 95% CI of $a$ | $r^2$ |
| --- | --- | --- | --- | --- | --- | --- |
| Whole plant | Mass | 0.647 | 0.584, 0.702 | 0.185 | 0.122, 0.258 | 0.898 |
|  | Surface area | 0.685 | 0.625, 0.738 | 0.0637 | 0.0487, 0.0807 | 0.892 |
| Roots | Mass | 0.712 | 0.641, 0.774 | 0.158 | 0.0913, 0.247 | 0.917 |
|  | Surface area | 0.631 | 0.588, 0.675 | 0.0288 | 0.0228, 0.0359 | 0.944 |
| Leaves | Mass | 1.09 | 0.977, 1.20 | 6.37 | 2.69, 13.9 | 0.792 |
|  | Surface area | 1.12 | 0.921, 1.27 | 0.598 | 0.203, 1.27 | 0.739 |
| Stems | Mass | 0.811 | 0.756, 0.875 | 0.292 | 0.187, 0.490 | 0.883 |
|  | Surface area | 0.972 | 0.896, 1.06 | 0.791 | 0.464, 1.54 | 0.879 |

In all regression, number of observation = 55, using RMA on log-log coordinates (for all cases, $P < 0.001$ ).

**Fig. 1.**
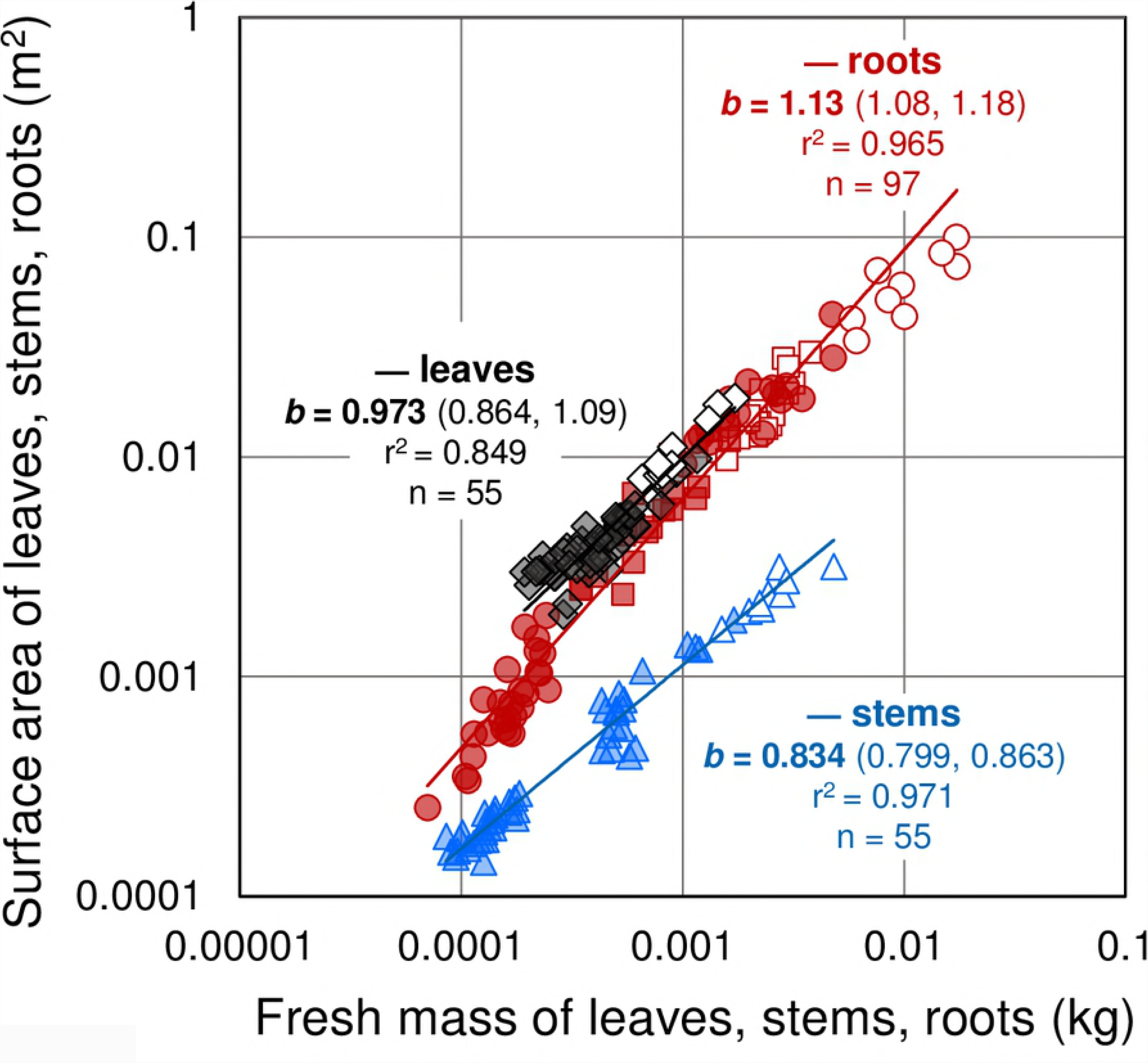
Relationship between whole-plant respiration and (a) whole-plant fresh mass and (b) whole-plant surface area. Each point is an individual seedling of current-year (filled, n = 46) and 1-year-old (open, n = 9). An exponent *b* (with 95% confidence interval) is the RMA slope fit to all points in each relationship.

### Respiration of organs

To see how organ-specific respiration contribute to the size-scaling of *R* shown in Fig. 1, we analyzed the size-scaling values for area in relation to mass (Fig. 2), and evaluated the relationships between respiration of roots, leaves, and stems (*R*_R_, *R*_L_, *R*_S_; μmol s^−1^) and their fresh mass (*M*_R_, *M*_L_, *M*_S_; kg) and surface area (*S*_R_, *S*_L_, *S*_S_; m^2^) separately, at the whole-organ level (Fig. 3, Table 2).

**Fig. 2.**
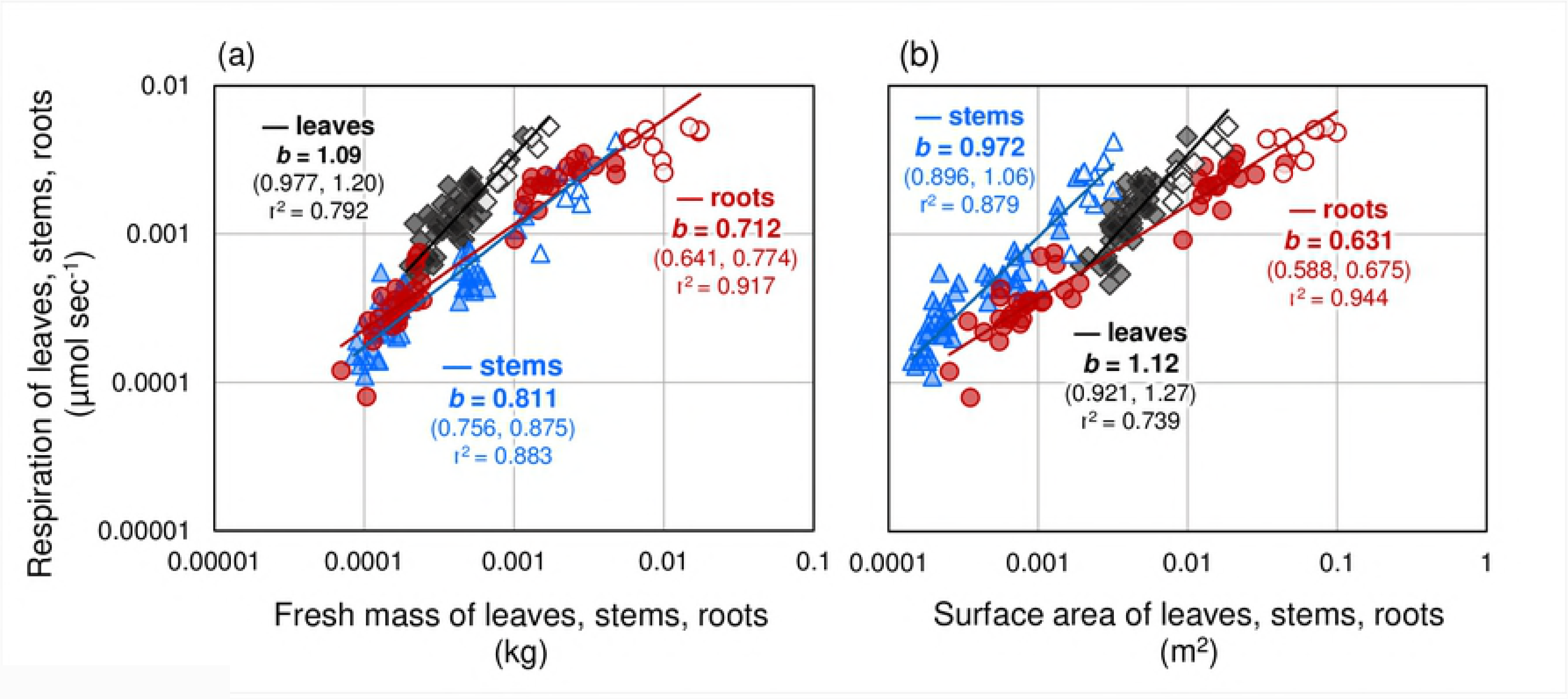
Relationships between surface area and fresh mass in plant organs at the whole-organ level. Each point is total leaves (diamonds, n = 55), stems (triangles, n = 55), and roots (circles and squares, n = 97) of current-year (filled) and 1-year-old seedlings (open). Roots depicted by squares (n = 42) were obtained from current-year seedlings (n = 16) and 1-year-old seedlings (n = 26) that were not used for respiration measurement. An exponent *b* (with 95% confidence interval) is the RMA slope of equation (1) fit to all points in each organ-specific relationship on log-log coordinates.

**Fig. 3.**
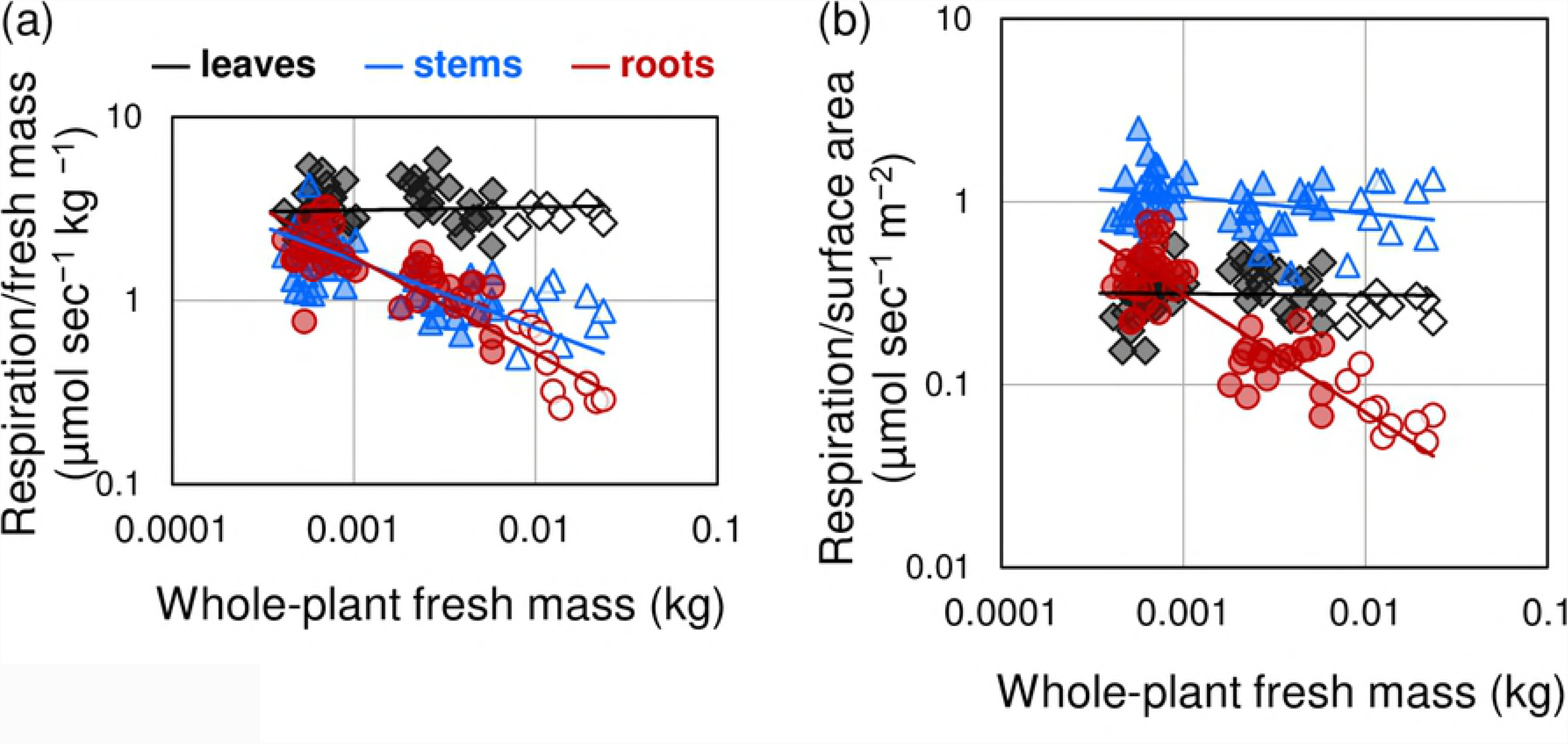
Relationships between respiration and (a) fresh mass and (b) surface area in plant organs. Each point is total leaves (diamonds), stems (triangles), and roots (circles) of the individual seedling in Fig. 1 of current-year (filled, n = 46) and 1-year-old (open, n = 9). An exponent *b* (with 95% confidence interval) is the RMA slope fit of equation (1) to all points (n = 55) in each relationship.

The roots had a 246-fold range in mass (*M*_R_) and 393-fold range in surface area (*S*_R_), revealing that *S*_R_–*M*_R_ relationship showed a significantly positive allometry, as shown in Fig. 2. On the other hand, root respiration (*R*_R_) varied within only 65.8-fold range, providing significantly low *b* values for the size-scaling of *R*_R_ to *M*_R_ (*b* = 0.712, 95% CI of *b*: 0.641 to 0.774, r^2^ = 0.917; Fig. 3a) and *R*_R_ to *S*_R_ (*b* = 0.631, 95% CI of *b*: 0.588 to 0.675, r^2^ = 0.944; Fig. 3b). Although the difference of *b* values was not significant, the exponent *b* of *R*_R_ to *S*_R_ was relatively lower than that of *R*_R_ to *M*_R_, indicating that the increase in *S*_R_ was more efficient than that in *M*_R_. This seems to represent an energetically efficient growth to effectively enhance water acquisition by increasing absorptive surface area with minimum carbon cost.

The leaves showed an 8.91-fold range of mass (*M*_L_) and 9.64-fold range of surface area (*S*_L_), providing nearly isometric scaling of *S*_L_*–M*_L_ relationship in Fig. 2. Further, the 11.6-fold range of leaf of respiration (*R*_L_) was close to that of *M*_L_ and *S*_L_. Consequently, we found that leaves had isometric scaling for both relationships of *R*_L_ to *M*_L_ (*b* = 1.09, 95% CI of *b*: 0.977 to 1.20, r^2^ = 0.792; Fig. 3a) and *R*_L_ to *S*_L_ (*b* = 1.12, 95% CI of *b*: 0.921 to 1.27, r^2^ = 0.739; Fig. 3b).

The stems had a 56.3-fold variation in mass (*M*_S_) and 22.2-fold range of surface area (*S*_S_), revealing a negative allometric scaling of the *S*_S_*–M*_S_ relationship in Fig. 2. Respiration (*R*_S_) showed a 38.5-fold range, which was smaller than the range of *M*_S_, but larger than that of *S*_S_. Consequently, the scaling of *R*_S_ to *M*_S_ was negatively allometric (*b* = 0.811, 95% CI of *b*: 0.756 to 0.875, r^2^ = 0.883; Fig. 3a), whereas the scaling of *R*_S_ to *S*_S_ was nearly isometric (*b* = 0.972, 95% CI of *b*: 0.896 to 1.06, r^2^ = 0.879; Fig. 3b).

### Respiration per unit mass and surface area of organs

We evaluated the respiration rate (energetic cost) per unit mass and surface area of organs, as related to total individual mass (*M*) on the log-log coordinates, fitting the relationships with a simple power function (1) (Fig. 4 and Table 3). For the roots, both *R*_R_/*M*_R_ (*b* = −0.456, 95% CI of *b*: −0.541 to −0.381, r^2^ = 0.751; Fig. 4a) and *R*_R_/*S*_R_ (*b* = −0.575, 95% CI of *b*: −0.642 to −0.497, r^2^ = 0.781; Fig. 4b) decreased with increasing *M*, resulting in the steepest change among the three organs. Conversely, for the leaves both *R*_L_/*M*_L_ (*b* = 0.00415, 95% CI of *b*: −0.0470 to 0.0516, r^2^ = 0. 000379; Fig. 4a) and *R*_L_/*S*_L_ (*b* = −0.00695, 95% CI of *b*: −0.0720 to 0.0511, r^2^ = 0.000846; Fig. 4b) were almost constant, regardless of *M*. Finally, for the stems *R*_S_/M _S_ declined with increasing *M* (*b* = −0.242, 95% CI of *b*: −0.309 to −0.167, r^2^ = 0.460; Fig. 4a), but *R*_S_/*S*_S_ was independent of *M* (*b* = −0.0644, 95% CI of *b*: −0.135 to 0.0216, r^2^ = 0.0511; Fig. 4b).

**Table 3.** Scaling of respiration per unit mass (µmol sec^−1^ kg ^−1^) and per unit surface area (µmol sec^−1^ m^−2^) of roots, leaves, and stems (n = 55) with whole-plant mass fitted by equation (1).

| Dependent variable | Exponent ($b$) | 95% CI of $b$ | Normalization constant ($a$) | 95% CI of $a$ | $r^2$ |
| --- | --- | --- | --- | --- | --- |
| $R_R/M_R$ | -0.456 | -0.541, -0.381 | 0.0708 | 0.0389, 0.109 | 0.751 |
| $R_R/S_R$ | -0.575 | -0.642, -0.497 | 0.00565 | 0.00380, 0.00930 | 0.781 |
| $R_L/M_L$ | 0.00415 | -0.0470, 0.0516 | 3.25 | 2.35, 4.24 | 0.000379 |
| $R_L/S_L$ | -0.00695 | -0.0720, 0.0511 | 0.300 | 0.200, 0.425 | 0.000846 |
| $R_S/M_S$ | -0.242 | -0.309, -0.167 | 0.285 | 0.189, 0.458 | 0.460 |
| $R_S/S_S$ | -0.0644 | -0.135, 0.0216 | 0.652 | 0.405, 1.12 | 0.0511 |

In all regression, number of observation = 55, using OLS on log-log coordinates.

**Fig. 4.**
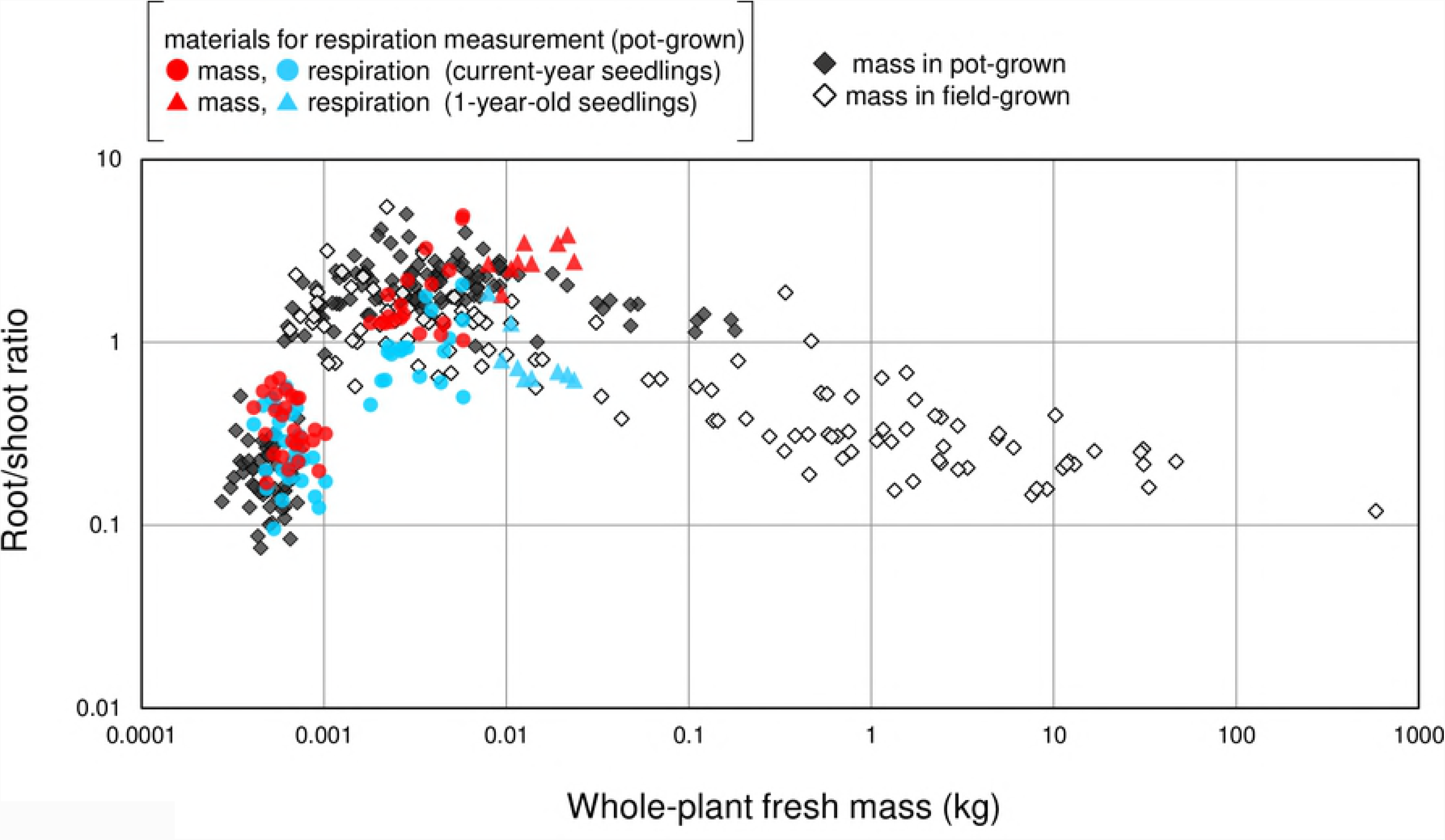
(a) Respiration/fresh mass and (b) respiration/surface area of plant organs in relation to whole-plant fresh mass. Each point is total leaves (diamonds), stems (triangles), and roots (circles) of the individual seedling in Fig. 1 of current-year (filled, n = 46) and 1-year-old (open, n = 9). Lines are OLS regression line of equation (1) fit to all points in each relationship, and details of the regression analysis are compiled in Table 3.

These results indicate that only roots show a significant decline in the energetic cost per unit surface area as well as per unit mass with increasing whole-plant mass. On the regression line, the *R*_R_/*S*_R_ declined about 90% and the *R*_R_/*M*_R_ declined about 84%, indicating that energetic cost per unit surface area declined more rapidly than that per unit mass.

### Partitioning of individual mass, surface area, and respiration to organs

The scaling of whole-plant respiration is determined by 1) the relative contribution of each organ to total mass and surface area, and 2) the organ-specific respiration per unit mass and surface area. Table 1 shows the maximum and minimum values of respiration, mass, and surface area of organs in the seedlings from current-year to 1-year old. It indicates that the proportion of roots to whole plant increased greatly both in mass and surface area with increasing *M*; the proportion of roots to whole plant in mass increased from approximately 17% (7.04/41.2) to 74% (1730/2350) and that in surface area increased from about 11% (2.54/23.6) to 82% (998/1210). Nonetheless, the increment of the proportion of root respiration was from 9.2% (0.800/8.70) to only 40% (52.6/131), revealing that the increment in the proportion of roots to whole plant is much more larger in mass and surface area than in energetic cost.

These results revealed that the similarity between the *R*–*M* (Fig. 1a) and *R*–*S* (Fig. 1b) relationships in their scaling exponents (Table 2) was largely due to the equally significant increase in the proportion of root mass and area. Hence, the combined effect of increasing root proportion and the decreasing mass- and area-specific respiration in the roots induced the negative allometry in the scaling of whole-plant respiration.

## Discussion

### Role of low-cost initial burst of root development

The significantly negative allometry of *R*_R_ to *S*_R_ (Fig. 3b) that provided the drastic decrease in *R*_R_/*S*_R_ (Fig. 4b) indicate that the root development in two years after germination is energetically efficient and effective for enhancing water and nutrient acquisition with minimum energy cost. The decrese in the energetic cost at root surface area (*R*_R_/*S*_R_) is probably mainly due to consumption of energy reserves in seeds. During the initial growth stage after the exhaustion of energy reserves, the photosynthetic performance remains lowest among whole life stages [29–31], and gradually increases accompanied by accumulation of nutrient with leaves thickening [29]. Therefore, the low-cost burst in root development seems to be reasonable and indispensable process for seedlings to induce the increase in photosynthesis. At the same time, this process would help to avoid water and nutrient deficit of tiny seedlings.

### Ontogenetic shift in root/shoot ratio from seedlings to mature trees

Our study suggests the need for further work at the whole-plant level up to mature trees, to clarify the role of low-cost rapid root development, beyond the initial seedling stage. Figure 5 depicts the relationship between root/shoot ratio in mass (*M*_R_/*M*_Shoot_ = *M*_R_/(*M*_L_ + *M*_S_)) and whole-plant mass of *F. crenata* from seedlings to mature trees (n = 346, compiled in S1 File), including the materials in this study (n = 55) and pot-(n = 178) and field-grown (n = 113) individuals from our prior work [10,32]. This figure also shows root/shoot ratio in respiration (*R*_R_/*R*_Shoot_ = *R*_R_/(*R*_L_ + *R*_S_)) for the 55 current-year and 1-year-old seedlings, and indicates that their *M*_R_/*M*_Shoot_ increases much greater than *R*_R_/*R*_Shoot_ with increasing *M*.

**Fig. 5.**
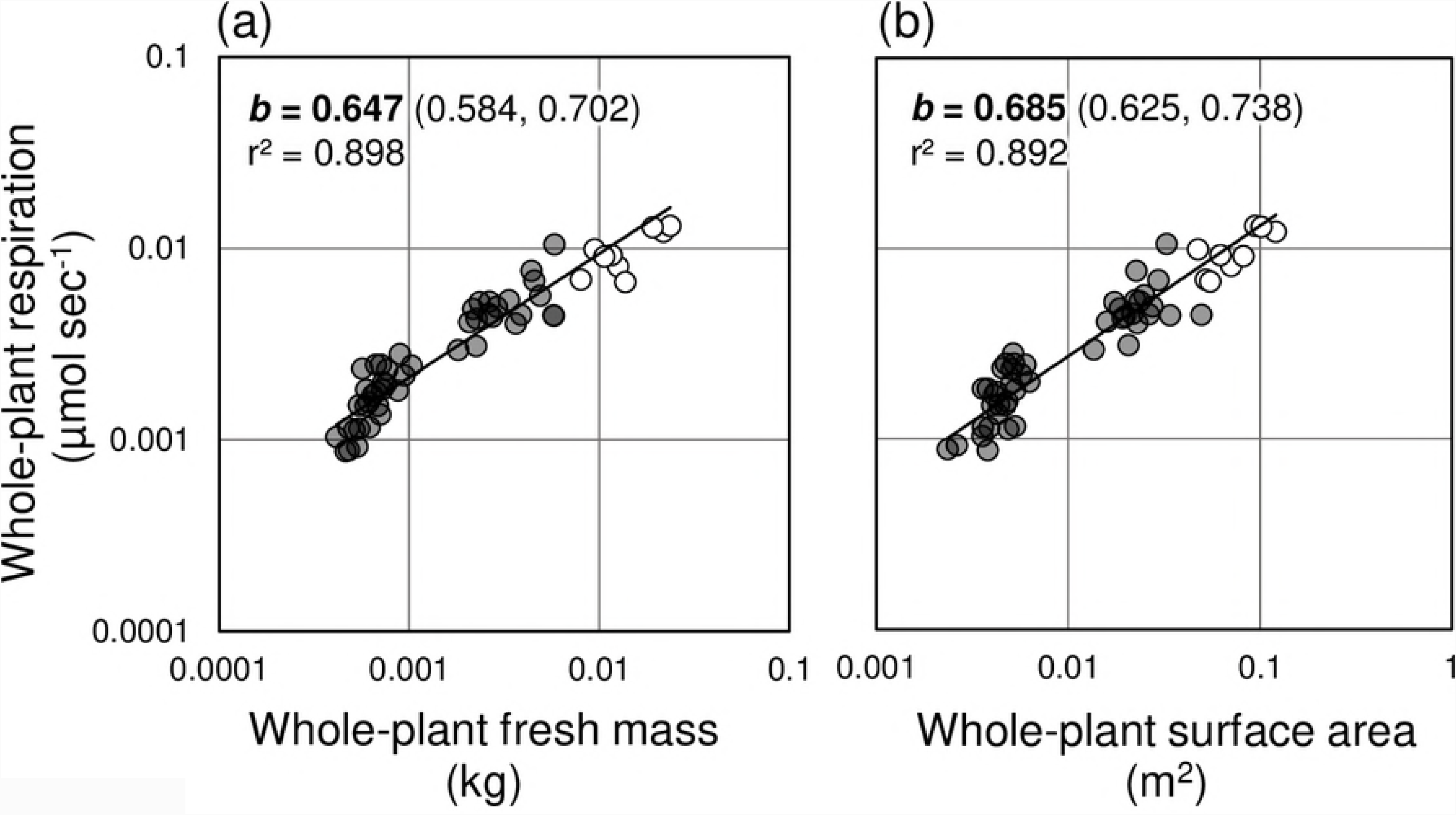
Plot of root/shoot ratios against whole-plant mass of *Fagus crenata* from germination to mature trees. The total plot number of the root/shoot ratio in mass is 346 (compiled in S1 File). Red and blue symbols represent the root/shoot ratio of the 55 seedlings in mass and respiration, respectively (circles; current-year seedlings; n = 46, triangles; 1-year-old seedlings; n = 9). Filled and open diamonds represent root/shoot ratio in mass of pot-grown (n = 178) and field-grown (n = 113) individuals, respectively that were obtained from our prior work.

As to *M*_R_/*M*_Shoot_, the rapid increment is specific of the seedling stage, but after that, it gradually declines with size (and presumably age) in both pot- and field-grown individuals. The decrease in *M*_R_/*M*_Shoot_ after the seedling stage should coincide with the gradual increase in photosynthetic performance during ontogenetic transition [29–31], entailing a decrease in *R*_R_/*R*_Shoot_. Since measurements were performed in healthy individuals, it is probable that the individuals that did not reach high *M*_R_/*M*_Shoot_ during the first year after germination had already died. Therefore, keeping low *M*_R_/*M*_Shoot_ during the initial growth stage may be one of the physiological reasons for death of seedlings in the bottleneck phase [22–25], and seedling survival under natural conditions is likely to depend on the rapid and low-cost development of roots at the individual level.

During the development from seeds to seedlings, the source that activates individual metabolism shift from chemical energy in seeds to current assimilation after initiation of photosynthesis [1,2,33]. This shift in the energy source may successively generate the low-cost burst in root development with shifting individual structure and function that would help to avoid water and nutrient deficit [11,17], and reduce mortality of seedlings.

### Importamce of lowering energetic cost for various individual organisms

Banavar et al. [34] suggested that plants and animals have reached equivalent energetic efficiencies through their independent evolution, using our prior data on whole-plant respiration from seedlings to giant trees [10]. Fundamentally, the studies on improvement of energetic efficiency have focused on animal locomotion and often suggested energy-saving mechanism as a strategy for effective resource acquisition [35–38]. On the other hand, although the resource acquisition is essential for all individual organisms including both plants and animals, very few studies have focused on energetic efficiency of terrestrial plants. Hence, whole-plant level empirical data on energetic cost in shoot and root would be important for us to obtain the understanding of physical and physiological constraints on metabolic scaling of terrestrial plants [10,39].

Plants are rooted in one place and they must acquire resources under continuously changing environment. Therefore, it is expected that plant metabolism must be flexible, relying on rapid adjustments in energetic efficiency. In this respect, the negative allometry of whole-plant respiration shown in Fig. 1 seems to indicate an improvement in flexibility for resource acquisition with body size, involving changes in the energy partitioning among organs. Therefore, the initial spurt and reduction in energetic cost in root development (shown in Figs. 2–4) could be considered as one of the underlying processes to effectively improve whole-plant energetic efficiency. In that case, it may be an energy-saving process of seedlings that is comparable to that in animal locomotion for resource acquisition [35–38]. In the present study, the measurement at the whole-plant level revealed the drastic reduction in energetic cost for rapid root development that would underpin the population dynamics and sustainability of the Northern Hemisphere forests that are dominated by *Fagus* trees [19–21]. The understanding of metabolic scaling of individual organisms would help to scale up the structure and function from organ level to ecosystem level [40–42].

## Acknowledgments

We thank T. Ichie and A. Iio for offering seed materials on this research. We also thank Y. Iiduka and D. Arai, staff of the University Forest, Faculty of Agriculture, Yamagata University, for assistance with our field work.

## Supporting information

**S1 File. Primary data.** Fresh mass, surface area, and respiration rate of total leaves, stems, and roots at the individual level.

